# Neurite growth kinetics regulation through hydrostatic pressure in a novel triangle-shaped neurofluidic system

**DOI:** 10.1101/2021.03.23.436675

**Authors:** B. G. C. Maisonneuve, A. Batut, C. Varela, J. Vieira, M. Gleyzes, J. Rontard, F. Larramendy, T. Honegger

**Author notes:** **Corresponding author:** Thibaut Honegger.

## Abstract

Microfluidic neuro-engineering design rules have been widely explored to create *in vitro* neural networks with the objective to replicate physiologically relevant structures of the brain. Several neurofluidic strategies have been reported to study the connectivity of neurons, either within a population or between two separated populations, through the control of the directionality of their neuronal projections. Yet, the *in vitro* regulation of the growth kinetics of those projections remains challenging. Here, we describe a new neurofluidic chip with a triangular design that allows the accurate monitoring of neurite growth kinetics in a neuronal culture. This device permits to measure the maximum achievable length of projecting neurites over time and to report variations in neurite length under several conditions. Our results show that, by applying positive or negative hydrostatic pressure to primary rat hippocampal neurons, neurite growth kinetics can be tuned. This work presents a pioneering approach for the precise characterization of neurite length dynamics within an *in vitro* minimalistic environment.

## Introduction

Mimicking the structure of neural circuits *in vitro* by using microfluidics^1^ technologies enables the efficient reconstruction of neuronal networks in comparison to conventional culture systems. Latter systems include Petri dishes^2,4^, where links between neurons are mainly created based on cell body proximity, and organotypic brain slices^3^, where neuronal networks are partially harmed because of the slicing process and might represent an incomplete portray of the brain’s neural circuitry. In comparison, microfluidic-based technologies are capable of confining and manipulating cell microenvironments^4^, allowing the *in vitro* synthesis of networks between different co-cultured neuronal cell types^5,6^. The rationalization of complex microfluidic designs is required to adapt neuronal network architectures into robust *in vitro* models. To date, most microfluidic chip designs applied to neuroscience research are based on the original model proposed by Taylor and colleagues in 2005, in which the connection between compartmentalized neuronal populations was achieved through guiding neurite outgrowth by using microchannels^2^. This system has the advantage to prevent the migration of cell bodies between culture channels while allowing only neuronal projections to pass through^2,6^. In addition, the longitude of those microchannels acts as a selectivity barrier for the exclusive passage of axons over dendrites^7,10^, and it also promotes the unidirectional growth of the neuronal projections from one compartment to the other. Myriad studies have focused their research on obtaining improved directional networks through the implementation of geometrical and dimensional modifications on asymmetric^8^ and rerouting axonal diodes^9^, the increase of microchannel number^10,24^ or the use of electrokinetic confinement^11,12^. Nevertheless, it is crucial to develop microfluidic systems to control not only neurite directionality, but also neurite outgrowth dynamics. Here, we introduce a neurofluidic chip with a novel triangle-shaped design that allows the exploration of neurite growth dynamics within asymmetric microchannels of various lengths. We observed that, when culturing primary rat hippocampal neurons in one compartment of our triangular microfluidic system, a maximum neurite elongation level is reached. Moreover, such neurite length can be regulated by applying different hydrostatic pressure (HP) gradients on the neuronal population. This work shows that, with this new device, it is possible to monitor the dynamics of neurite outgrowth *in vitro,* providing a suitable platform for studying the development of connections among compartmentalized neuronal populations.

## Materials and Methods

### Device design and masks fabrication

The device is a two-layered compartmentalized chip with two separated chambers connected by a set of asymmetric microchannels, represented as a right triangular shape with angles of 53° and 37° (Figure 1A). The design of the bottom layer (3-μm thick) contains 20-μm interspaced asymmetric microchannels from 5 μm to 7 μm width and lengths ranging from 225 to 10725 μm (inset Figure 1A). The microchannels are narrow enough to only permit the passage of neurites to the second microfluidic compartment in a directional way, whereas neuronal bodies stay inside the first one (Figure 1B, 1C). To avoid any experimental issues that might be caused by the fluidic resistance of the microchannels, both plating channels of the device were fluidically isolated, similarly to previous publications^17^. The upper layer presents identical geometry for both compartments but does not include the microchannel patterns. This mask is used to increase the final height of the loading chambers to 100 μm.

**Figure 1:**
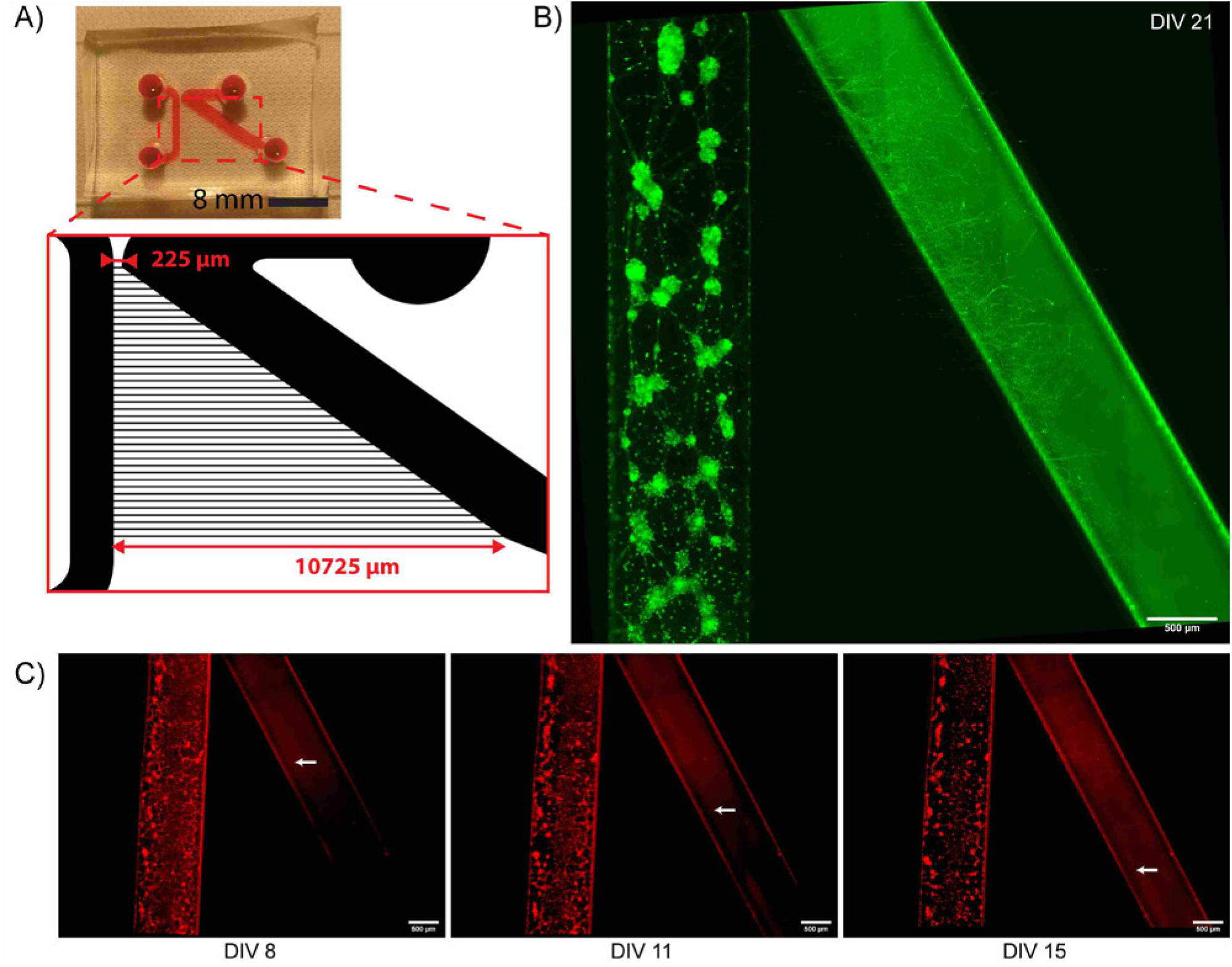
***(A)*** General structure of the triangle-shaped neurofluidic system filled with rhodamine B. *Inset:* Schematic representation of the different lengths of the microchannels and the position of the plating compartments. ***(B)*** 21 DIV rat hippocampal neurons were stained with Vybrant™ DiO Cell-Labeling Solution for live-cell imaging. The image shows how the growing neurites exit the microchannels of various longitudes to reach the neuron-free channel. Scale bar indicates 500 μm. ***(C)*** Selection of live-cell images at different culture time points, where rat hippocampal neurons were stained with Vybrant™ DiD Cell-Labeling Solution. The white arrows indicate the longest microchannel through which neurites extended at those specific time points. Scale bars indicate 500 μm.

Respective masks for the designed layers were fabricated using a homemade mask-less UV photolithography system^13^, or purchased on transparent films (Selba SA, CH).

### SU-8 mold fabrication

This procedure was adapted from one of our previously published protocols^13^. The mold was fabricated using a silicon wafer as substrate. An adhesion promoter (Omnicoat, MicroChem, USA) was spin-coated onto the wafer at 300 rpm for 30 sec, before letting it bake at 200 °C for 1 min. SU8-2002 (MicroChem, USA) was spin-coated onto the wafer at 1000 rpm for 30 sec, followed by a soft bake step (1 min at 65 °C, 2 min at 95 °C, and 1 min at 65 °C). Using the mask corresponding to the bottom layer of the microfluidic chip, the wafer was exposed to an appropriate dose of UV light (90 mJ/cm^2^) using a MJB4 mask aligner (Süss MicroTech, Germany). After exposure, another soft bake step was done (1 min at 65 °C, 2 min at 95 °C, and 1 min at 65 °C) before developing the wafer in SU8 developer (MicroChem, USA). The wafer was finally rinsed with isopropanol. SU8-2050 was then spin-coated onto the same wafer at 1667 rpm for 30 sec, followed by another bake step (5 min at 65 °C, 16 min at 95 °C, and 5 min at 65 °C). The second mask was aligned with the SU8 features already onto the wafer using alignment marks, before exposing the wafer to UV light (230 mJ/cm2). A post-exposure bake step was performed (4 min at 65 °C, 9 min at 95 °C, and 2 min at 65 °C) and the wafer was developed in SU8 developer again, before being washed with isopropanol. A final hard bake step (150 °C for 15 min) was done.

### PDMS microfluidic chip fabrication

This procedure was adapted from one of our previously published protocols^13^. Briefly, the wafer was first silanized using vapors of a silanizing agent (trichloro(1H,1H,2H,2H-perfluorooctyl)silane) in a desiccator for 30 min. Polydimethylsiloxane (PDMS) prepolymer (Sylgard 184, Dow Corning, USA) was prepared by mixing the base and the curing agent at a ratio of 10:1. The mixture was degassed, poured onto the wafer and cured at 80 °C for 40 min. The PDMS layer was cut to the required size, peeled off the wafer, and the inlet/outlet zones were punched out. The device and a clean microscope glass slide were then plasma treated using a plasma cleaner (Harrick Plasma, USA) before their assembly. The device was then sprayed and filled with a solution of 70% of ethanol and brought into a sterile environment. The ethanol was washed away three times using sterile distilled water and exposed to UV light for 30 min.

The plating channels within the chambers of the microfluidic device were then coated with 0.1 mg/mL poly-L-lysine (Sigma Aldrich, USA) for 24 hours in an incubator. The channels were then rinsed three times with Hank’s Balanced Salt Solution (HBSS) (Life Technology, Thermo Fisher Scientific Inc., USA) buffered with 10 mM 4-(2-hydroxyethyl)-1-piperazineethanesulfonic acid (HEPES) (Life Technology, Thermo Fisher Scientific Inc., USA) and coated with 20 μg/mL laminin (Sigma Aldrich, USA) for 2 hours. The coated channels were washed again with HBSS and filled three times with hippocampal culture medium composed of Neurobasal-B27 (Life Technology, Thermo Fisher Scientific Inc., USA) containing 2 mM glutamine and 100 U/mL penicillin/streptomycin (Life Technology, Thermo Fisher Scientific Inc., USA). The microfluidic chips were placed in an incubator until use.

### Neuron preparation and seeding

According to previously published procedures^13^, hippocampi were harvested from E18 OFA rats (Charles River Laboratories) and kept in ice-cold HBSS buffered with 10mM HEPES (pH 7.3). The tissue was digested for 30 min using 2 mL of HEPES-buffered HBSS containing 20 U/ml of papain (Worthington Biochem., USA), 1 mM EDTA (PanReac AppliChem) and 1 mM L-cysteine (Sigma Aldrich, USA). Then, the tissue was rinsed three times with 8 mL of hippocampal culture medium. The cells were gently triturated in 1 mL of hippocampal culture medium, counted with a hemocytometer, and flowed into the device. The cells were maintained under incubation conditions (37 °C, 5% CO2, and 80% humidity) until use.

Before seeding, the reservoirs of the microfluidic chip were emptied without removing the media from the channels. Through one inlet reservoir, 4 μL of high density (> 8×10^6^ cells/mL) dissociated neuron solution were placed near the entrance of the channel. The chip was returned to the incubator for 5 min in order to let the neurons adhere on the coated surface, and the seeding process was repeated three times to achieve a high cell density. Finally, each input and output reservoirs of the device were filled with hippocampal culture medium, and chips were returned to the incubator.

All animal work was approved by the CEA, CNRS Ethics Committee of Animal Care, and abided by institutional and national guidelines. Experiments performed at NETRI were approved by regional authorities for animal welfare (DDHS Agreement SPA-2019-19).

### Neuron culture in device

The neurons seeded within the device were cultured up to 21 days *in vitro* (DIV), and the culture media was renewed three times per week. Cultured cells were subjected to three HP conditions, where the difference of static pressure at the air/fluid interface was controlled between both microfluidic compartments through the addition of extra medium at the inlet and outlet of the chamber: (i) no addition of extra volume of medium (0 μL), (ii) addition of 40 μL of extra medium, and (iii) addition of 80 μL of extra medium. To study the effect of negative pressure, those HP conditions were applied to the opposite chamber of the device, where no neurons where seeded. The difference in volume induces a height variation of the free surface and, therefore, a difference in pressure. Taking into consideration that the inlet and outlet cylindrical openings of the chambers have a radius of 2 mm, the addition of ±40 μL and ±80 μL simulates a pressure difference of ±30 Pa and ±60 Pa, respectively. The induced HP was calculated from the applied variation of volume by using Bernoulli’s principle.

### Live-cell imaging

For neurite outgrowth visualization in the triangle-shaped microfluidic system, neurons in culture were imaged at several time points until reaching 21 DIV. For each time point, cells were stained using the Vybrant™ Multicolor Cell-Labeling Kit (V22889, Thermo Fisher Scientific Inc., USA).

For live-cell visualization in the standard microfluidic system, 21 DIV neurons were stained also using the Vybrant™ Multicolor Cell-Labeling Kit Vybrant Cell-Labeling Solutions, incubating cells in one channel with 5μM DiO Labeling Solution, and cells in the other channel with 5μM DiD Labeling Solution.

Live-cell imaging was performed using an AxiObserver Z1 Microscope (Zeiss, Germany) and micromanager^14,15^. The obtained images were stitched using the FiJi software^16^. The staining process was done on each compartment separately, without inducing hydrostatic pressure changes during the entire procedure.

### Neurite length measurements

Neurite length variations were determined over time using an automated algorithm implemented in MATLAB 2020b (MATLAB 2020b, The MathWorks Inc., USA) by sequentially binarizing the images using an appropriate thresholding and measuring the distance from the microchannel openings on the seeded compartment and the end of the growing neurite. The maximum length of the neurites was calculated as the length of the longest microchannel the neuronal projections went through. A home-made ImageJ plugin was developed to boost image analysis throughput. Source code can be provided upon request.

### Statistical analysis

Statistical analysis was determined using one-way ANOVA with a post-hoc Tukey’s test. The statistical software used for the analysis was Origin Pro 8.0 (OriginLab Corporation, USA). All reported points were replicated at least three times in independent experiments.

## Results and discussion

### Triangle-shape design discriminates neurite extension

In order to trigger variations on neurite length kinetics, we built a microfluidic chip that contained two separated compartments connected by microchannels of different lengths. The combination of this atypical set of microchannels with both culture channels represents the shape of a right triangle on the device (Figure 1A). Such triangular design allows discretizing the maximum neurite extension (projecting from a neuronal population) by strategically applying a length increase of 33 μm per microchannel. We examined the filtering capabilities of the structural design of our device by only seeding primary rat hippocampal neurons on one of the compartments (Figure 1B), similarly to previous protocols for design characterization^8,9^. Seeded neurons were maintained in culture until reaching 21 days *in vitro* (DIV) to compare, in terms of neurite development, early and late culture stages^18,19^.

Fluorescent images acquired at different time points indicate that, the more time seeded neurons are maintained in culture, the more projections of passage manage to exit longer microchannels (Figure 1C, SI 2), suggesting that this microfluidic design can be used to classify neurite elongation stages depending on the neuronal maturation status.

### Neurite elongation is arrested after 12 days in culture

Considering the observed reliability of the triangle-shaped system, we next studied the evolution of neurite elongation process in order to identify a maximal neurite growth length. To do so, we monitored neurite length across several DIV within our device while applying different HP gradients. The longest microchannel from where neurites exited to the empty plating channel at defined culture time points was pinpointed, which corresponded to a defined neurite projection length. Results indicate that, when no HP was used (0 μL condition), the growth of the neurites reached a plateau after 12 DIV, defining a neurite length of 2.4 mm that rested constant over time (Figure 2A) Such value can be represented as the maximum growth length that neurites projecting from hippocampal neurons can attain when going through microchannels, which is one of the limiting factors when designing *in vitro* neural networks.

**Figure 2:**
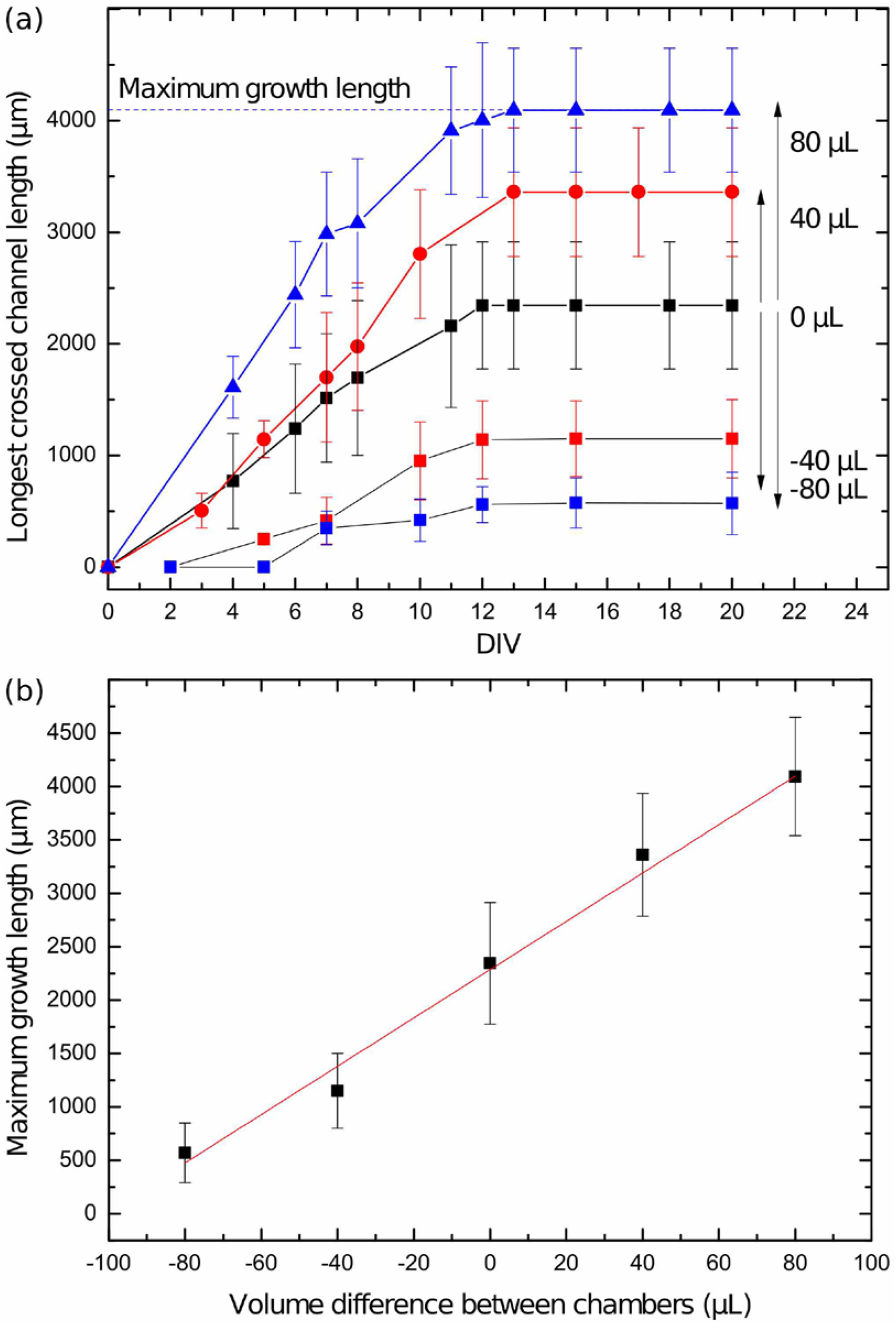
***(A)*** Plot indicating the length of the longest microchannel in which neurites projected (μm), in relation to the age of the neuronal culture (DIV), for several hydrostatic conditions: no applied pressure (0 μL of extra medium), positive pressure (extra 40 μL or 80 μL on seeded channel), and negative pressure (extra 40 μL or 80 μL on cell-free channel). The maximum growth length is represented for the 80 μL hydrostatic condition. ***(B)*** Graph plotting the determined maximum growth length values according to the several volume difference between both compartments. The red line represents the linear interpolation. All data indicated as mean ± SD; (n = 3).

Our data might be linked to previous findings on *in vitro* neuronal network development and matureness. Several studies have shown that the maturation degree of *in vitro* neuronal cultures increases drastically between 12 DIV^19,20^ and 14 DIV^21,22^, moment in which neurite arborization reached stability. During this time span, the electrical activity of the cultures exhibits increased firing rates, constant burst patterns, and also highly synchronized periods of high frequency activity^20,22^. Moreover, 14 DIV and older neuronal cultures show a significant increase in synapse density per area unit compared to younger culture stages, supporting the idea of a mature connected network^19,20^.

Based on these observations and our results, one could speculate that neurons in culture undergo three different transformation stages: (i) an initial expansion stage, in which neurites grow during the first two weeks of culture; (ii) a following connection stage, where neurites attain their maximum elongation point and lead to a structurally stable network before the neurons reach 14 DIV; and (iii) a final functional stage, in which the established neurite connections trigger synapse formation and plasticity, leading to a mature and functional network in neurons older than 14 DIV. However, to verify our hypothesis, network functionality between 12 DIV and 21 DIV should be assessed by examining synaptic composition and electrical properties of the neurons in culture.

### Neurite length cap can be extended using pressure gradients

In order to overpass the maximal length limitation without using any chemical cues, we investigated the influence of HP gradients on neurite outgrowth dynamics. Such gradients were created by adding 40 μL or 80 μL of extra hippocampal culture medium at both the inlet and outlet reservoirs of the channel where the neurons were seeded. We observed an increase of maximum neurite length from 2.4 mm (0 μL condition) to 3.4 mm and 4.1 mm for the 40 μL and 80 μL conditions, respectively, indicating that the presence of positive HP on the seeded channel has a slight effect on the extension of the neurites (Figure 2A). Remarkably, the growth of the neurites seems to hit a plateau region after approximately 12 DIV, independently of the three HP conditions. This observation indicates that the application of a small pressure gradient does not seem to modify the time frame of the transformation stages that were previously described. Hence, the use of HP might only have an influence on the maximum length of the neuronal projections. As shown in previous studies, improving the directionality of growing projections results in larger distances covered by those projections^23^.

Although it still remains unclear how HP is able to enhance neurite orientation and growth, it is possible that the use of HP regulates the directionality of developing neurites by applying weight onto the growth cones, and hence forcing them to align with the flow streamlines. Nevertheless, this speculation falls beyond the scope of this present work.

### Neurite extension can be mitigated using negative pressure

Inversely to our previous experiments, we have also explored the capacity of HP to confine neurite growth into one compartment using negative pressure (*i.e.*, by applying a difference on HP from the cell-free channel to the one where cells were seeded). Our results indicate a limited extension of the neurites through the microchannels over time (Figure 2A), suggesting that the application of negative pressure might decelerate neurite growth kinetics. Interestingly, we still report a plateau region reached by the growing neurites after 12 DIV, which is consistent with our previous observations and represents the maximum growth length of the neurites within microchannels.

When examining the obtained maximum growth length values in relation to the various HP conditions applied between plating channels, we can determine a linear correlation between the difference in volume and the maximal length values for each pressure condition (R=0.981) (Figure 2B). These data indicate a direct first-degree mechanical correlation between growth cone forces and hydrodynamical forces at stakes within the microchannels.

### Positive and negative HP might influence neurite growth orientation

We took advantage of our observations to investigate the impact of HP for improving structural directionality between two populations of neurons. For this experiment, primary rat hippocampal neurons were seeded in both plating channels of a standard compartmentalized microfluidic chip and maintained in culture with a constant application of 80 μL extra volume of medium on the right-sided compartment during 14 DIV (+80 μL). As shown on SI 1, most identified neurites (80%, n=254 microchannels) grew from the compartment where the HP was positive towards the other compartment. In regard of this result, it is tempting to hypothesize that positive HP might affect the guidance of neurite outgrowth to a specific direction within microchannels.

Even though the *in vitro* regulation of neurite orientation falls beyond the scope of this work, the perspective of studying HP as a new microfluidic method to control structural directionality of neuronal networks should be considered. Despite the fact that such technique might not be able to compete with other highly efficient unidirectional approaches^8,9^, HP-induced neurite growth could be used in the future as a simple and easy method to orientate neural cultures in neurofluidic chips without the need to use chemical compounds.

### Conclusions and perspectives

There is an urgent need to generate physiologically relevant neural circuits within an *in vitro* minimalistic environment. Thanks to our unique and innovative neurofluidic design, we can efficiently monitor and measure neurite outgrowth kinetics, as well as quantifying the maximum neurite length according to the application of varying pressure gradients.

We believe that our work apports a new method to effectively modify and enhance the maximum growth length of neuronal projections by only using hydrostatic pressure rather than artificial chemical cues. In this regard, the knowledge of the maximal length that neurites can reach within microchannels will be useful when intending to design improved *in vitro* neural circuits with specific connectivity patterns.

In conclusion, this study apports valuable neurofluidic design strategies for the investigation of neurite length dynamics, which is essential for the *in vitro* analysis of the integrity and complexity of the brain. To further strengthen the efficacy of those strategies, future work should focus using such methodology to evaluate the effect of pharmacological compounds on neurite elongation and orientation on maturing and mature human curlure as well as proving the impact of hydrostatic pressure on neural architecture manipulation.

## Author Contributions

BM, JR and TH designed the research rational; BM, JV and FL designed and fabricated the devices; BM, AB, CV, MG carried out the animal and in vitro experiments. BM, JR, MG and TH performed data analysis and co-wrote the manuscript.

## Acknowledgements

We thank Dr. Clara Berenguer Escuder (CBE Science Writing) for assisting with the structural organization of this manuscript, and for the revision and editing of the original draft.

## Supplementary Information

**SI 1:** Rat hippocampal neurons were seeded on both compartments of a standard microfluidic chip, which are connected by a set of microchannels of 450 μm-long. 21 DIV cells were stained using the Vybrant™ Multicolor Cell-Labeling Kit for live-cell imaging, incubating cells on the left channel with DiO solution (green), and cells on the right channel with DiD Solution (red). The neurons were maintained in culture under constant hydrostatic pressure from the left (green) compartment to the right (red) compartment. Scale bar indicates 200 μm.

**SI 2:** Time-lapse video of primary rat hippocampal neuron culture within the triangle-shaped neurofluidic system.

