## Supplementary figures and images for "Neurite growth kinetics regulation through hydrostatic pressure in a novel triangle-shaped neurofluidic system"

### Figure SI 1

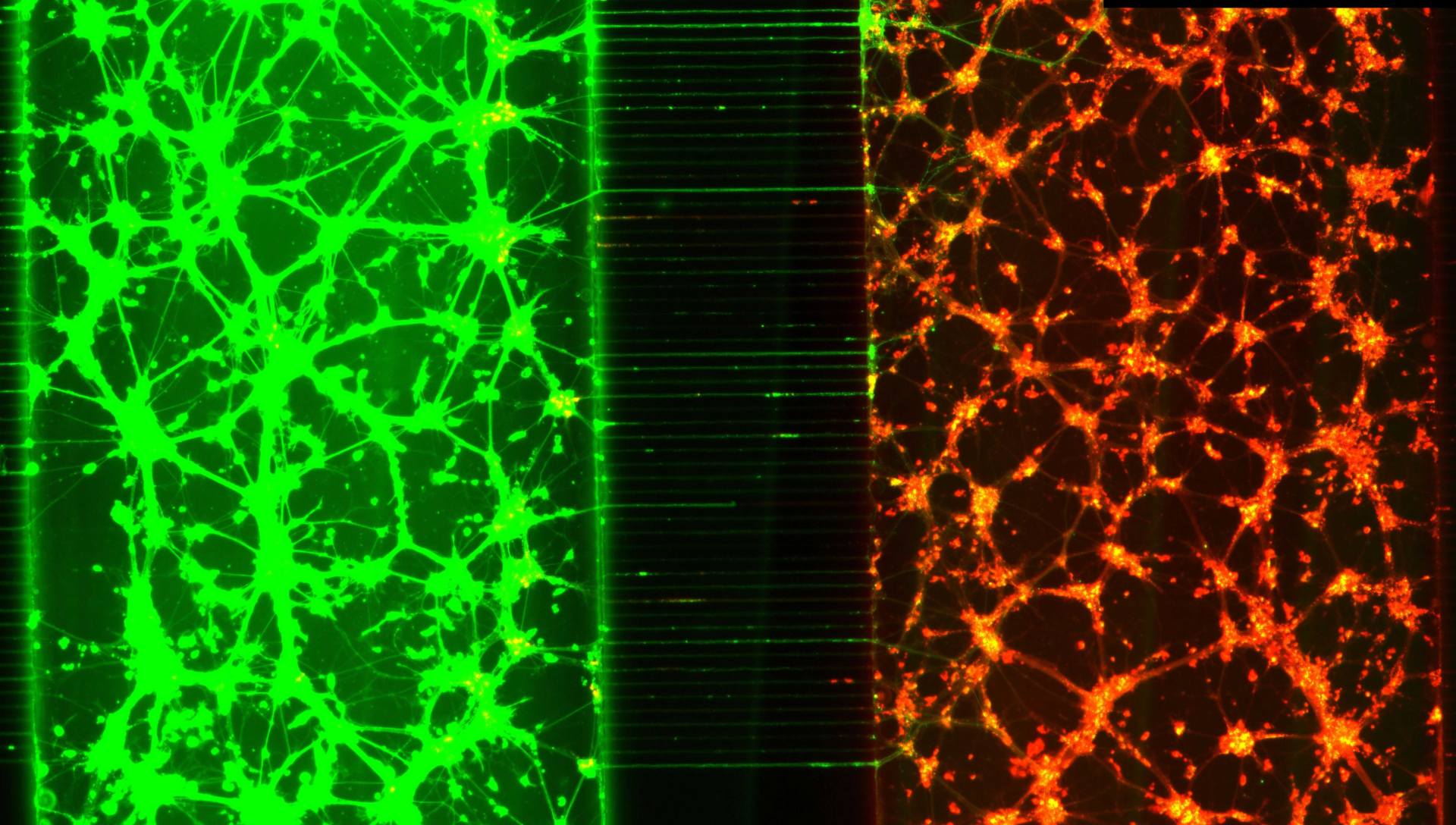
